# Label-free differentiation of functional zones in mature mouse placenta using micro-Raman imaging

**DOI:** 10.1101/2023.07.21.550049

**Authors:** Arda Inanc, Nayce Ilayda Bektas, Ibrahim Kecoglu, Ugur Parlatan, Begum Durkut, Melike Ucak, Mehmet Burcin Unlu, Ciler Celik-Ozenci

## Abstract

In histopathology, it is highly crucial to have chemical and structural information about tissues. Additionally, the segmentation of zones within a tissue plays an important role in investigating the functions of these regions for better diagnosis and treatment. The placenta plays an important role in embryonic and fetal development and in the diagnosis of some diseases associated with its dysfunction. This study provides a label-free approach to obtain the images of mature mouse placenta together with the chemical differences between the tissue compartments using Raman spectroscopy. To generate the Raman images, spectra of placental tissue were collected using a custom-built optical setup. The pre-processed spectra were analyzed using statistical and machine learning methods to acquire the Raman maps. We found that the placental regions called decidua and the labyrinth zone are biochemically distinct from the junctional zone. Comparison and evaluation of the Raman maps with histological images of the placental tissue were performed by a histologist and they are found to be in agreement. The results of this study show that Raman spectroscopy offers the possibility of label-free monitoring of the placental tissue from mature mice while revealing important structural information about the zones at the same time.

## Introduction

Raman spectroscopy is one of the vibrational spectroscopy techniques that offers a non-invasive and label-free detection of samples at the molecular level. In addition to various gases and liquids, it is widely used to probe the molecular structure of solid samples such as art and archaeological objects [1], crystals [2], and especially for biological samples including cells and tissues [3]. The identification of the chemical compounds in the biological samples allows researchers to diagnose or monitor the progression of certain medical conditions by comparing the biomarkers specific to the specimen found in Raman spectroscopy experiments [4–6]. Raman spectra can be collected over a range of areas by scanning the sample plane, which is commonly called Raman imaging. The biochemical information obtained in this way can be used for segmentation [7–10].

Tissue segmentation and zone assessment are critical aspects of biomedical research. This is usually achieved by one or a combination of the modern staining methods described in the literature [11], which are inherently used for in-vitro studies, and by the evaluation of a histopathologist. However, in recent in-vitro studies, tissues are segmented using artificial intelligence (AI) on the images of Hematoxylin and Eosin (H&E) stained tissues [12–14]. As Raman spectroscopy is a non-invasive and label-free technique, unlike staining methods such as H&E, it has great potential for in-vivo imaging [15–17] and is gaining interest for use in tissue imaging and segmentation.

The placenta plays a vital role in the development of the mammalian embryo, regulating the exchange of gases, nutrients, and waste products between the mother and the fetus. It is a source of hormones linked to pregnancy and also supports the immune defense system of the fetus. Despite its crucial role, little is known about the molecular causes of human pregnancy diseases, which arise from disorders in the placenta. As the mouse and human placentas are structurally similar and express many of the same genes that control placental growth and functioning, the mouse placenta is a particularly effective model for understanding the human placenta [18–20]. The biochemical origins of some human pregnancy complications, such as pre-eclampsia, first-trimester miscarriage, intrauterine growth restriction (IUGR), and preterm labor can be elucidated by studying placental development in mice. It is therefore of great importance to carry out experimental research on mouse placental tissue in order to gain a better understanding of the human placenta.

The mature mouse placenta is structurally and functionally composed of three distinct compartments, each with a unique role in supporting the developing fetus: the maternal decidua (D), the junctional zone (JZ), and the labyrinth zone (LZ). The maternal D, the outermost layer of the placenta, lies close to the mother’s uterine wall, providing structural support, and allowing the placenta to attach to the uterus. It contains uterine decidual cells, maternal vasculature, glycogen trophoblast (GlyT) cells, and spiral artery-associated trophoblast giant cells (SpA-TGC). The JZ, the middle layer of the placenta, acts as a source of hormones, growth factors, and energy required for normal placental and embryonic growth and is composed mainly of the parietal TGC (p-TGC) layer, spongiotrophoblast (SpT), and nonmigratory GlyT cells. GlyT cells function as an energy reservoir by storing glycogen and as a source of insulin-like growth factor II (IGF-II) [21]. Meanwhile, SpT cells and TGCs are the primary sources of hormones and growth factors [22]. The LZ is the largest component of the placenta, and it is essential for proper placental function, but its organization and function are intricate and multifaceted.

Despite the structural differences highlighted above, the functional placental villous units of mice and humans are remarkably similar [22, 23]. In both cases, maternal blood flows through sinuses lined by trophoblast cells, specifically the SynT cells, representing the primary site for nutrient and gas exchange. Therefore, proper placental function is critical for normal developmental progression during intrauterine development.

In this study, the micro-Raman spectroscopy technique was employed to distinguish the three main zones of mature mouse placenta by analyzing the biochemical information obtained from these zones without labeling. Upon completion of the analysis of the collected Raman spectra, the regions corresponding to D and LZ were found to be biochemically and visually distinct from JZ, where the biochemical differences between the zones were proved to be consistent. Furthermore, it was discovered that imaging the paraffinized placental tissues for a selected wavenumber in the Raman spectrum made the morphology of the tissue easily accessible without additional algorithms and techniques for digital deparaffinization, and the overlapping effect of paraffin on the tissue lipid is demonstrated through image analysis. Thus, Raman spectroscopy is shown to be an excellent candidate to replace other conventional staining techniques such as H&E staining for morphological and biochemical characterization of tissues.

## Results

### Raman clustering maps and histological assessment

Placental tissues were obtained from mature mouse placentas and prepared according to the procedures described in subsections Sample collection and preparation and Tissue processing and Hematoxylin and Eosin staining. Prepared placental tissue samples from three different mice were scanned using the custom-built micro-Raman spectroscopy setup described in subsection Raman microspectroscopy and scanning. After an image edge detection was applied to eliminate the non-tissue regions, the 3D Raman spectral data were preprocessed, where the substrate spectrum was also subtracted from the tissue spectra at this stage. All the details about edge detection and preprocessing are given in subsection Edge detection and preprocessing. Principal component analysis (PCA) and k-means clustering with three clusters were then applied to the normalized spectra, which provided the clustered Raman maps. For comparison with the Raman clustering maps, the scanned tissues were also H&E stained and imaged as described in subsections Tissue processing and Hematoxylin and Eosin staining and Histology and image analysis.

For each mouse, H&E stained tissue images are given in Fig 1a, where the three main zones, LZ, JZ, and D, were labeled and outlined by a histologist. The labeled H&E stained images of the tissues were cropped and aligned with the Raman clustering maps for a better visual comparison of the zones. The Raman clustering maps are shown in Fig 1b. As can be seen, the Raman clustering maps of the tissues have three different zones represented by three different colors. The corresponding Raman spectra are given in Fig 1c with the same color as the groups given in the cluster maps, while the standard deviations of the average spectra of the groups are given in shaded [24].

**Fig 1.**
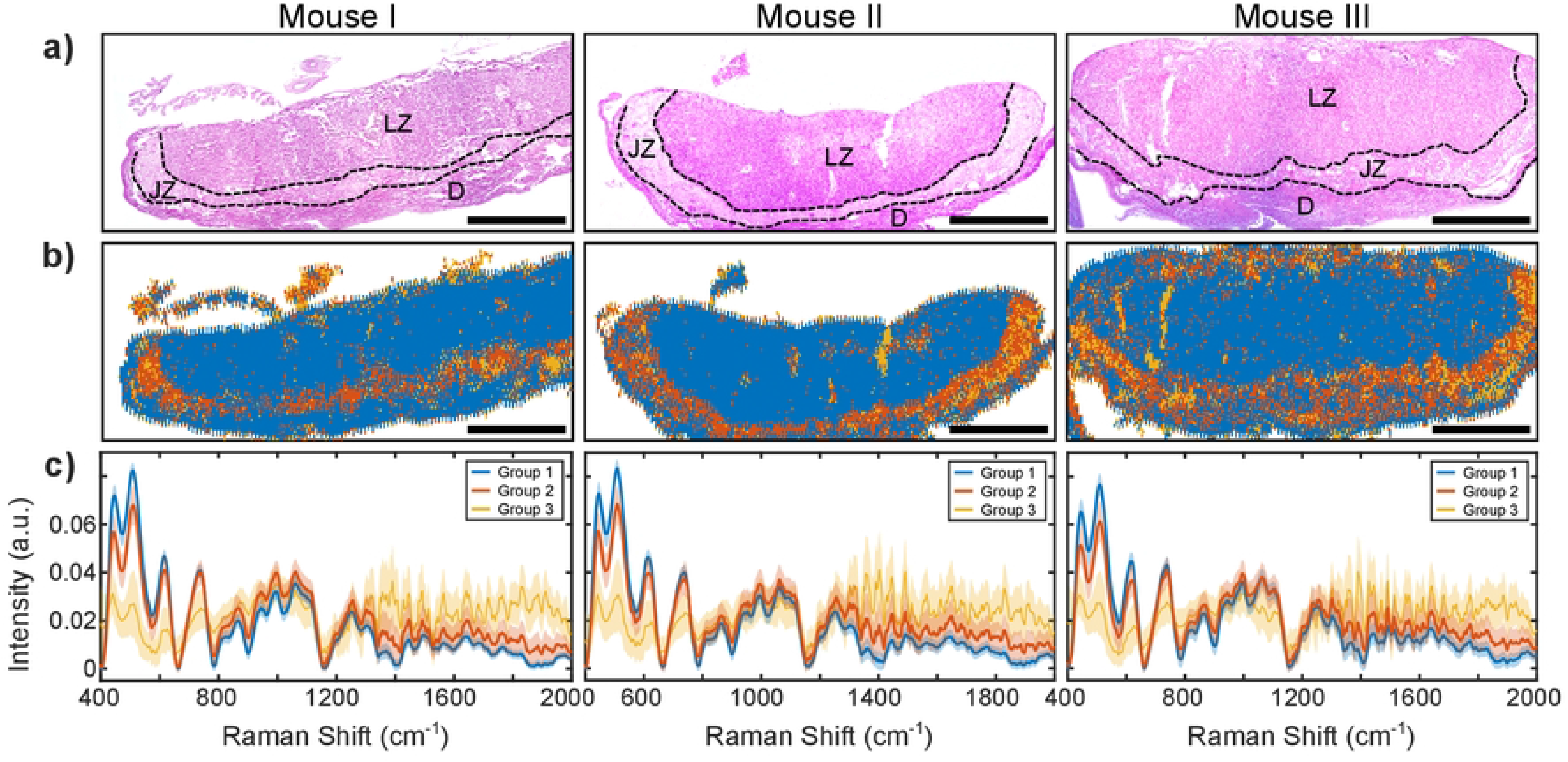
a) H&E stained images of placental tissues, where the three main zones, LZ, JZ, and D, are identified and outlined by a histologist. b) Raman clustering maps of the tissues, where the three main zones are obtained by k-means clustering. The scale bars are in 1 *mm*. c) Corresponding Raman intensities of the zones represented in the maps. Standard deviations of the Raman intensities are shown in the same color but shaded.

From the comparison of the Raman clustering maps in Fig 1b with the corresponding H&E stained tissue images in Fig 1a, it was observed that there is a considerable spatial alignment between group 1 map and the zones D and LZ, while group 2 map was found to be correlated mostly with the JZ. Before further analysis of the zones, an inference was made about group 3 in the Raman clustering maps. Empty regions were observed in the H&E staining images in Fig 1a. Since these empty regions mostly coincide with group 3 map (yellow areas) on the corresponding Raman maps in Fig 1b, it was concluded that group 3 may originate from the glass substrate. The spectra of this group given in Fig 1c, also fluctuate as a noisy signal with a large standard deviation. A noisy spectrum is expected for a region of the substrate because the substrate spectrum is subtracted during the polynomial fitting in the algorithm as described in subsection Edge detection and preprocessing. It was therefore concluded that the main spectral contribution to group 3 for all three placentas came from the glass substrate beneath the tissues and this group was not included in the further analysis as the filtered background regions around the tissues were not included in the analyzed data. As a result, the zones of the tissues were clustered into two groups, with the group 2 map (JZ) acting as a partition between the two zones within the map of group 1, separating D and LZ. In light of the histological findings, it is apparent that the two results are indeed compatible. This observation has been conclusively validated by the esteemed histologist. Moreover, the identified major zones mirror those reported in the referenced papers, which further substantiates our conclusions [25–27].

### Analysis of Raman spectra and tentative assignments

As the spectral decomposition of groups 1 (D + LZ) and 2 (JZ) for each tissue sample was found to be very similar and the conclusions drawn were the same, the Raman clustering map, the spectra of the groups, the PCA scatter plot for the first two PCs and the loadings are given in Fig 2a, b, c, and d for one mouse only. The resulting graphs for the remaining mice are given in Fig S1 of the S1 File. The Raman intensities of the groups and the loading plots, shown in Fig 2b and Fig 2d, respectively, are plotted with a common axis so as to mark the wavenumbers of the tentative assignments in the fingerprint region on both figures. Three loading plots for the first three principal components (PC) are shown with the corresponding total explained variances (TEV) given in parentheses.

**Fig 2.**
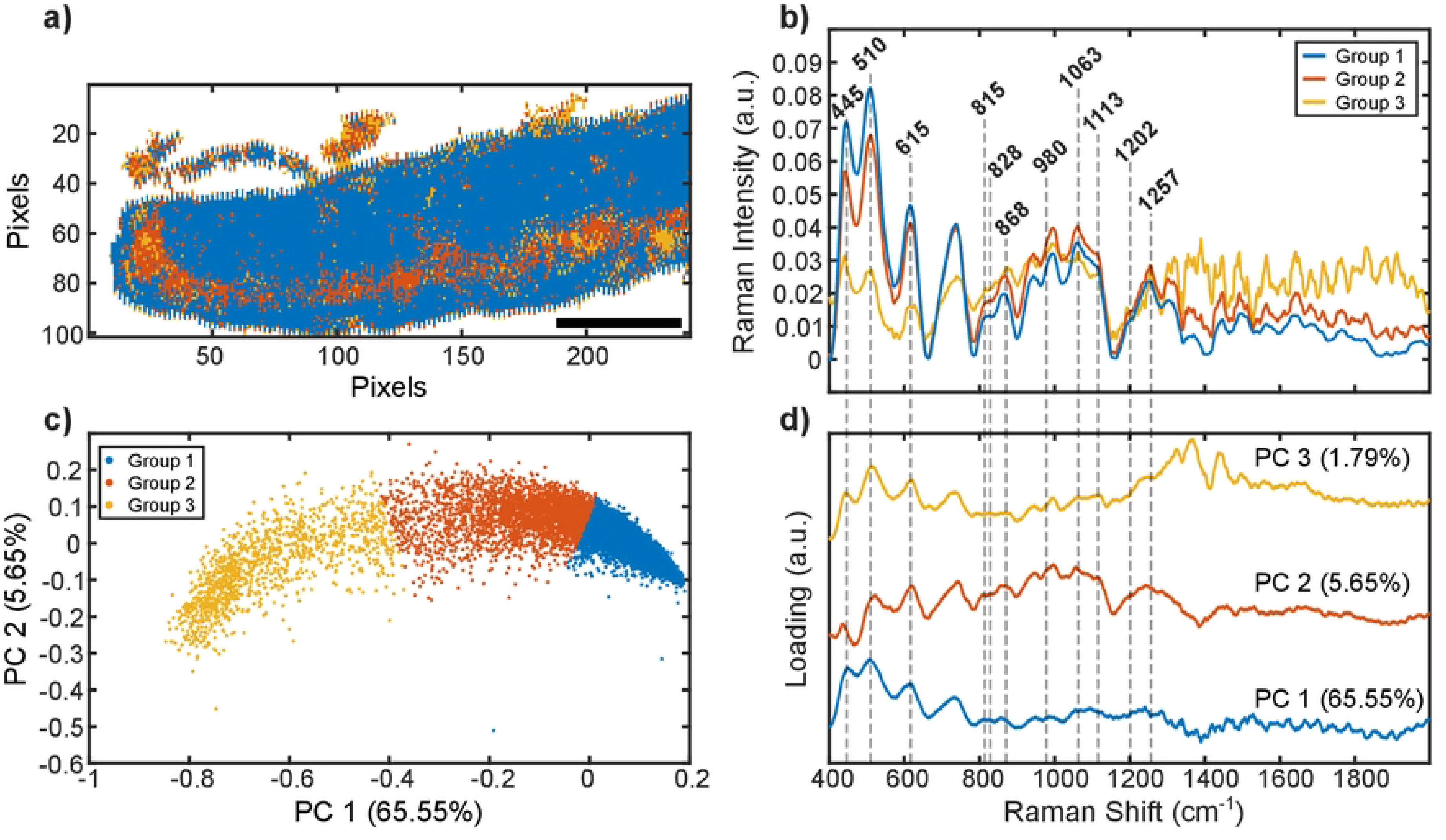
a) Raman clustering map of the placental tissue from the first mouse. The scale bar is in 1 *mm*. Here, the group 1 map was found to be associated with the placental region LZ and D, and is shown in blue, while the group 2 map was found to be largely coincided with JZ and represented in red. As described in the text, it was deduced that group 3 mostly represents the empty regions within the tissue boundaries b) Raman intensities of the groups after PCA and k-means clustering together with the tentatively assigned wavenumbers for the fingerprint regions. c) Scatter plot of the PCA scores for the first two PC. The percentages of the TEV are given in the parentheses next to the axis labels. d) Loading plots for the first three PCs. In figures b and d, the wavenumbers for the tentative assignments are given on a common Raman shift axis.

For the indicated Raman wavenumbers given in Fig 2, a tentative assignment table containing the vibrational modes, average normalized Raman intensities with their standard deviation, and the references for the assignments is provided in Table 1 [28]. Among the assigned wavenumbers, the bands at 445 *cm^−^*^1^, 510 *cm^−^*^1^, and 615 *cm^−^*^1^ were found to be larger in intensity in the average spectrum of group 1 than in group 2, while the reverse is true for the remaining wavenumbers. Regarding the corresponding tentative assignments indicated in Table 1, collagen, Cysteine, cholesterol ester, and Thiocyanate were found to be more abundant in group 1, which was spatially associated with D and LZ, whereas proline, hydroxyproline, glucose-related molecules, lipids, and Amide III were found to be more abundant in group 2, which was spatially associated with JZ of the placental tissue. A table showing the normalized Raman intensity of the other two mice at the assigned wavenumbers is given in the S1 File, Table S1.

**Table 1.**
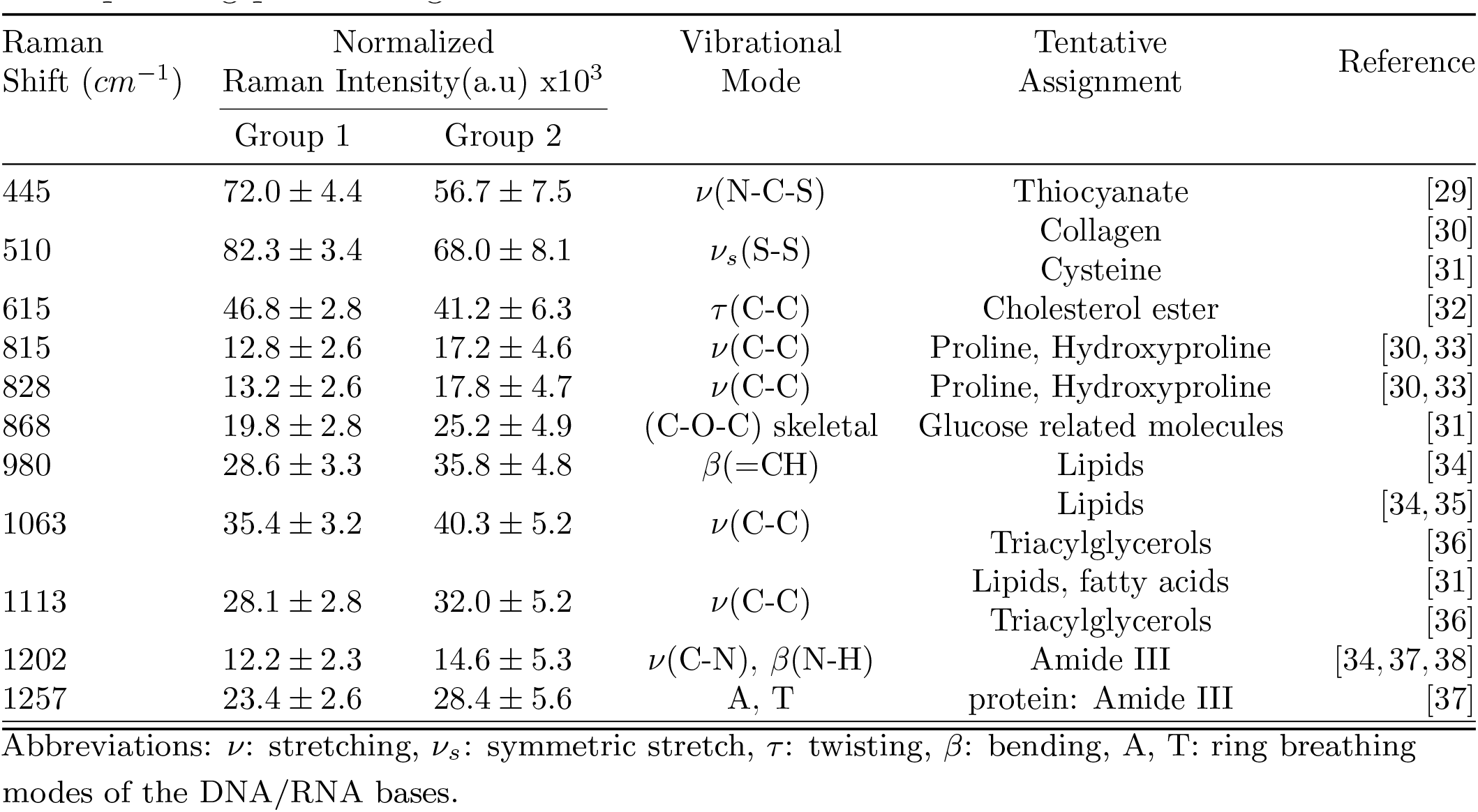
The tentative assignment table for the peaks marked in Fig 2b and Fig 2d is given below. For group 1 and group 2, average normalized Raman intensities of the corresponding peaks are given with their standard deviation.

In order to better distinguish between groups 1 and 2, violin plots for a few chosen wavenumbers are shown in Fig 3 in addition to the Raman clustering maps, the mean, and the standard deviation of the clustered groups. These two groups were subjected to a quantitative analysis using the Wilcoxon rank sum test with Bonferroni-Holm correction in MATLAB, and it was discovered that they were distinct with 99.9% confidence (*p <* 0.001) for all assigned wavenumbers in all three tissue samples. Violin plots at the six chosen wavenumbers for the other two mice are given in Fig S2 of the S1 File.

**Fig 3.**
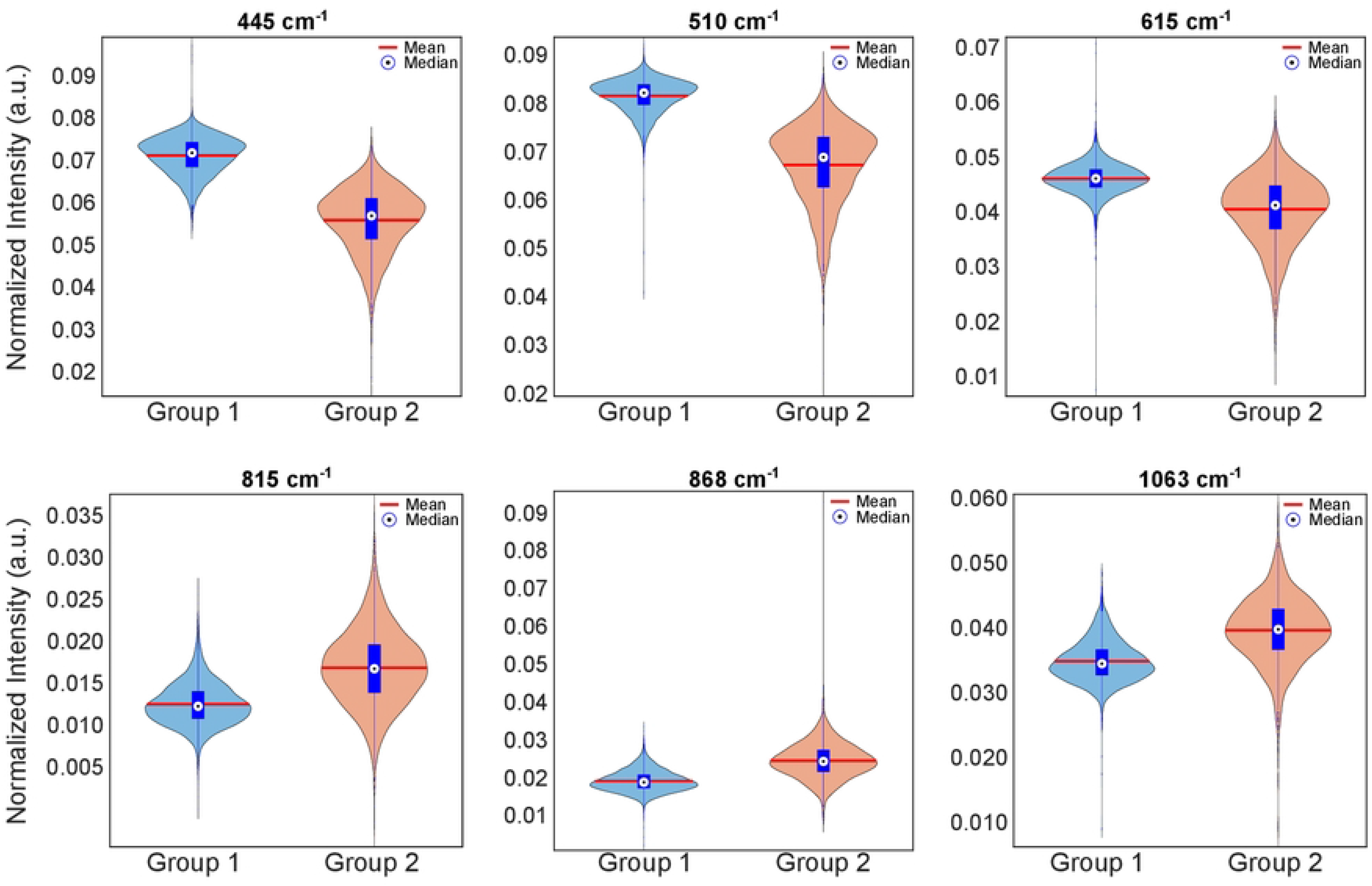
Violin plots for some selected wavenumbers showing the distribution of data within group 1 and group 2 found in clustering maps. Using the Wilcoxon rank sum test with Bonferroni-Holm correction in MATLAB, the null hypothesis that the two groups are the same was rejected at the 0.1% significance level (*p <* 0.001) for all three tissue samples. In the violin plots, the blue boxes represent the interquartile range covering the 25*^th^* and 75*^th^* percentiles of the data within the groups.

Taking into account all the above analyses, it was found that the two groups in the Raman clustering maps correspond to the three main zones of the mature mouse placenta. Histological assessments confirmed that the group 2 map spatially matches with JZ and divides the group 1 map into two regions corresponding to D and LZ. The label-free identification of the zones was achieved thanks to the diversity in the biochemical composition of the zones obtained by the Raman spectroscopy technique. The biochemicals that differentiate the groups, obtained from the analysis of the Raman spectra, are indicated in the tentative assignment table in Table 1. These differences between the groups were found to be statistically significant and are discussed in detail in section Discussion in terms of the importance of the tentative assignments for the placental zones.

### Imaging of placental morphology without deparaffinization

In this section, it is shown that the morphology of mature mouse placental tissue can be revealed without any chemical and digital deparaffinization. For deparaffinized tissue samples, the wavenumber 510 *cm^−^*^1^ was shown to be the most dominant peak in the Raman spectra contributing most to the differentiation of the zones in the PCA analysis. For paraffinized samples, 510 *cm^−^*^1^ was also found to be the most expressed peak on the tissues. In addition, among all the Raman images of automatically found peaks, 510 *cm^−^*^1^ was found to be the most promising peak that reveals the morphology of the tissues, where the Raman images at this wavenumber and the H&E stained images were found to be mostly consistent with each other. Thus, the wavenumber 510 *cm^−^*^1^ was used to image paraffin-embedded placental tissues and as a ground truth for the comparison of the images at different wavenumbers.

Paraffin is known to overlap with lipid content in tissues particularly between the wavenumbers 1000 *cm^−^*^1^ and 1700 *cm^−^*^1^ [39]. To show the effect of paraffin in masking the lipid content on the images, which was mostly found in the JZ, two other prominent wavenumbers in the lipid bands of the spectrum were also selected, namely 1060 *cm^−^*^1^ and 1120 *cm^−^*^1^. For mouse 1, the Raman images at these three wavenumbers with and without deparaffinization are given in Fig 4a together with the mean spectra before and after deparaffinization in Fig 4b.

**Fig 4.**
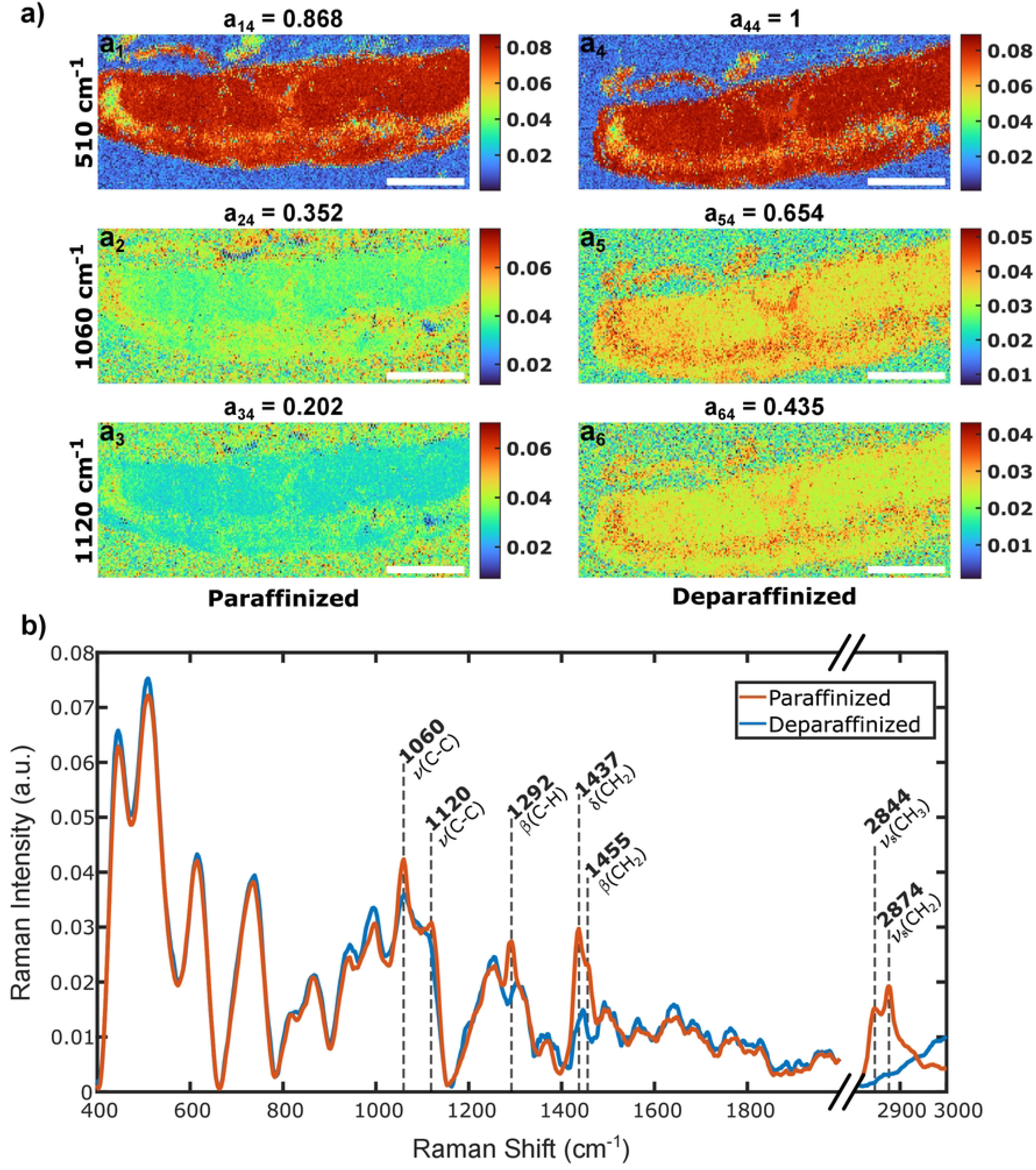
a) Raman images of placental tissue from mouse 1 at three different wavenumbers with and without deparaffinization. The values at the top of each image indicate the correlation coefficient calculated between the selected pairs. The scale bars are in 1 *mm* b) Average spectrum of the tissue before and after the deparaffinization. Abbreviations: *ν*: stretching, *ν_s_*: symmetric stretch, *δ*: deformation, *β*: bending.

The peaks in the spectrum of the paraffinized tissue that are more dominant than the peaks in the spectrum of the deparaffinized tissue were examined and the wavenumbers were found to be 1060 *cm^−^*^1^, 1120 *cm^−^*^1^, 1292 *cm^−^*^1^, 1437 *cm^−^*^1^, 1455 *cm^−^*^1^, 2844 *cm^−^*^1^, 2874 *cm^−^*^1^. These wavenumbers and their corresponding vibrational modes are shown in Fig 4b. Similar to the literature, the wavenumbers marked in the plot were found to be due to paraffin. A block of paraffin formed in the fixation process was measured separately and reported in the S1 File, Fig S3a [40, 41]. In order to compare all the Raman images for structural similarity, a geometric transformation including only translation and rotation is computed between the paraffinized and deparaffinized images at wavenumber 510 *cm^−^*^1^. In this way, the spatial offset between the two measurements was eliminated by applying the calculated transformation to all the paraffinized tissue images, and a correlation was calculated as described in subsection Machine learning and statistical analysis. The calculated correlation coefficients are given at the top of each image in Fig 4a, while the image pair being compared is indicated by the subscripts to the left of the coefficients. Since the image of the deparaffinized tissue scans at wavenumber 510 *cm^−^*^1^ was chosen as the ground truth, the Raman images at 1060 and 1120 *cm^−^*^1^ for both paraffinized and deparaffinized scans were compared with the deparaffinized tissue image at 510 *cm^−^*^1^. For the paraffinized tissue of mouse 1, the correlation coefficients between the images in Fig 4a_2_-a_4_ and Fig 4a_3_-a_4_ were calculated to be 0.352 and 0.202, respectively, indicating weak correlations. On the other hand, for the deparaffinized tissue, the correlation coefficients between the images in Fig 4a_5_-a_4_ and Fig 4a_6_-a_4_ were calculated to be 0.654 and 0.435, implying strong and moderate correlations, respectively. The correlation coefficient between the images in Fig 4a_1_-a_4_ was found to be 0.868, corresponding to a very strong correlation, while the correlation of the image in Fig 4a_1_ with itself provided the expected value of 1. The correlation strengths are defined at the end of the subsection Machine learning and statistical analysis.

The consistency and the importance of the biochemical findings in the Raman spectra analyses in relation to the placental zones are discussed in the next section.

## Discussion

Research on imaging of placental tissue sections dates back to 1897 [42], where a staining method is used for labeling. Very recent studies have reviewed the methods of imaging placental tissue, reflecting the importance of the topic [43–45]. Some of the prominent modalities used in placental imaging are ultrasound imaging [46, 47] magnetic resonance imaging (MRI) [48–50], micro-computed tomography (micro-CT) [51, 52], transmission electron microscopy and scanning electron microscopy [53–56], confocal fluorescence microscopy [57, 58], photo-acoustic imaging [59, 60], and immunohistochemistry [61, 62]. They have become to be used as ground truth for newly discovered methodologies. However, some of these methods are deprived of providing biochemical information about the tissue, despite the fact that they provide 3D in-vivo images. On the other hand, some of the remaining methods are not suitable for in-vivo imaging, while others require labeling. In this work, the three major zones of the adult mouse placenta were distinguished using the micro-Raman spectroscopy technique by examining the biochemical data collected from these zones without labeling.

The following subsections provide details of the comparison between the Raman spectroscopy results, whether the tentative assignments in Table 1 were found to be lower or higher relative to the zones, and the compatibility of existing knowledge about these zones.

### Junctional Zone

Even though JZ remains less understood, it is known to be responsible for hormone secretion, and the production of growth factors and cytokines necessary for normal placentation. The JZ provides energetic (glycogen), hormonal, and physical support to ensure proper placentation and pregnancy development. It has been shown that an increase or insufficiency in GlyT cells, and therefore total glycogen, can lead to IUGR [22, 63–65]. Glycogen is also thought to act as a reserve, storing energy and supplying nutrients to the placenta and embryo [22]. As the JZ is filled with GlyT cells, the absence of this structure or these cells can lead to severe birth defects caused by a lack of important energy reserves and hormone production during late gestation [22, 66]. In our study, glucose-related molecules such as monosaccharides (beta-fructose), disaccharides (Sucrose), polysaccharides, amylose, and amylopectin were successfully identified and the corresponding peak was found to be higher in JZ compared to D and LZ. In addition, proline, which plays a role in protein folding and stabilizing the structure of collagen molecules, and hydroxyproline, which is critical for the stability and strength of collagen fibers, were found to be higher in JZ compared to D and LZ. Taken together, these suggest the unique structure of JZ as a physical support to ensure that Raman spectroscopy can distinguish correct placentation. Amide III is a useful tool for studying protein conformational changes, folding, and structural stability of collagen, which is important for understanding the mechanical properties and physiological functions of connective tissues. It was found that the Raman bands corresponding to Amide III were higher in JZ than in D and LZ.

The placenta secretes many hormones into the maternal circulation, which modulates the mother’s physiology and thus transfers oxygen and nutrients to the fetus for growth. Lipids are also the building blocks of many hormones, including cholesterol-derived steroid hormones, progesterone, and estrogen. These hormones, secreted by the placenta, modulate most of the mother’s systems throughout pregnancy. They also regulate the production of other hormones, such as prolactin and placental lactogens, which in turn, may contribute to physiological changes in the mother as well. It was also found that the Raman intensities in the tentatively assigned lipid bands were higher in JZ, indicating the contribution of the known function of these molecules in hormone production.

### Decidua and Labyrinth Zone

Many of the most common placental lesions in the mouse are found in the LZ, the primary site of nutrient and gas diffusion [67, 68]. The LZ provides a circulatory pathway in which maternal and fetal blood are in close proximity, and insults to its architecture disrupt oxygen and nutrient exchange, leading to a definitive impact on fetal growth and, therefore, the placenta [69]. A prominent presence of cholesterol esters was found in the LZ and D regions of the placenta by Raman spectroscopy. It populates the hydrophobic core of circulating lipoproteins, delivering cholesterol and fatty acids to organs, in this case, to the fetus and/or the mother. Cholesterol esters can serve as a reservoir of cholesterol that can be mobilized for fetal development. Maternal plasma cholesterol would be obtained from external sources, specifically, the maternal circulation via the placenta, transported across trophoblasts, and effluxed or secreted into the fetal circulation and serves as a precursor for progesterone synthesis, and is essential for early fetal development.

The placental extracellular matrix (ECM), which is part of the establishment of materno-fetal interaction, not only contributes to structural support but also regulates cellular signaling-modulating processes such as proliferation and motility [70]. In our experiments, collagen was found to be higher in D and LZ compared to JZ. Different collagen types observed in the placenta have been described to be involved in fibril-forming (Col1a1), basement membrane (Col4a1 and Col4a2), beaded filament-forming (Col6a1, Col6a2a, and Col6a3), anchoring fibril-forming (Col4a1 and Col4a2), and multiplexing (Col18a1) collagens [71]. The presence of collagen types on the placenta attests to the maintenance of a three-dimensional architecture, where the thicker ones are supportive, and the thinner ones complement the ECM mesh structure, maintaining the labyrinthine capillary network and other structures [22, 72]. The cysteine was also found to be high in D and LZ. Cysteine is a non-essential amino acid required for protein production and other metabolic functions, and is important for collagen production. Cysteine metabolism is closely linked to the maintenance of redox status and protection against free radical oxidation, homocysteine elimination, and detoxification [73]. Overall, the distinctive presence of these proteins in D and LZ compared to JZ is comparable to the known function of these placental compartments in mice. Finally, a high level of thiocyanate was found in D and LZ compared to JZ. Thiocyanate has a role as a human metabolite. Thiocyanate functions in host defense as part of the secreted lactoperoxidase microbicidal pathway.

### Tissue morphology and biochemistry suppressed by paraffin

In tissue-based research, the fixation of a tissue is a prerequisite to prevent denaturation, and the use of formalin-fixed paraffin preserved (FFPP) tissues is a widely accepted procedure in this regard. However, paraffin produces strong Raman peaks in the spectrum that usually interfere with the fingerprint regions of interest and prevent the expression of biochemicals in these regions. There are recent papers in the literature that have studied the effect of paraffin wax on human breast tissue [74, 75]. Since Raman scattered light is negatively affected by paraffin and chemical removal of it can cause various other problems [76], there are papers that have developed digital deparaffinization algorithms and techniques to reveal the masked Raman spectral information by eliminating the effect of paraffin [39, 41, 77, 78]. In this study, it was found that imaging the paraffinized placental tissues for a chosen Raman wavenumber made the morphology of the tissue easily accessible without the need for additional algorithms and techniques for digital deparaffinization. The results obtained in subsection Imaging of placental morphology without deparaffinization show that the presence of paraffin blocked the realization of the tissue morphology, but it became visible after deparaffinization for both wavenumbers at 1060 *cm^−^*^1^ and 1120 *cm^−^*^1^. In particular, the expression of the zone corresponding to JZ was found to be increased compared to the other regions of the tissue after deparaffinization. The correlation coefficient between the images at 510 *cm^−^*^1^ was calculated to be 0.868, indicating a very strong correlation. The definition of the strength of association for correlation coefficient values is given in Table S2 of the S1 File. It was also observed from the images that the morphology of the tissue at 510 *cm^−^*^1^ is similar for both paraffinized and deparaffinized tissue scans. Thus, the images and the calculated correlation coefficients between them demonstrate that the morphological information of the tissue can be accessed by Raman imaging of placental tissue at 510 *cm^−^*^1^. As the chosen wavenumbers 1060 *cm^−^*^1^ and 1120 *cm^−^*^1^ are indicators of the lipid content in the tissue, it was also observed that the paraffin prevents the lipid expression, especially in JZ. This was confirmed by visual inspection of the intensity distribution on the images before and after deparaffinization and by the calculated correlation coefficient between the images. The results for the other two mice were found to be similar to those of the first, except that the last mouse placenta had increased expression of the other two zones as well at the specified wavenumbers when deparaffinized. The results for the other two mice are given in Fig S4 of the S1 File.

### Limitations and future work

In this study, tissues are paraffinized and sectioned at 5 *µm* thickness for histology, requiring a deparaffinization protocol prior to the Raman measurements, which may also get rid of the lipid contents of the tissue. The small thickness of the tissue results in less attenuation of the laser reflection and fluorescence from the glass. This, in turn, obscures the regions of the Raman spectrum obtained despite the use of a substrate removal algorithm. For the analysis of the pre-processed data, PCA and k-means clustering were applied in this project since the computational burden is less compared to the other dimension reduction and classification algorithms. However, the mapping from a higher to a lower dimension in PCA is a linear operation, which misses non-linearities that can be found in Raman spectra. As a classifier, the k-means algorithm alone cannot capture the relationship between classes that are related to other parameters, such as the spectral variances, because it only computes the mean of the spectra for classification. Also, the k-means algorithm suffers from the reproducibility of the results due to the selection of the starting points for the centroids, and spectra can move to another close cluster after an iteration [79]. As far as the imaging in the experiments is considered, the image area for each tissue cannot be known exactly prior to a scan. Therefore, a common fixed area was chosen to efficiently use the time at the expense of missing some parts of the tissues.

For the tissue sectioning and substrate problems, a snap-frozen thick section of around 20 *µm* can be collected and measured on CaF_2_ or cheaper metal substrates for high contrast spectral acquisition. In our laboratory, we are currently undertaking research to explore and contrast the limitations posed by tissue sectioning methods and substrate selections. Specifically, the study involves more cost-effective metal substrates for high-contrast spectral acquisition, and deparaffinized tissues that are sectioned at 5 *µ*m for histological analysis. By analyzing these procedures, we aim to develop a more optimized approach that minimizes such limitations, potentially improving the accuracy and reproducibility of Raman spectroscopic investigations in biological tissues. For preprocessing, an autoencoder as a deep learning network can additionally be used to incorporate the non-linearities and to denoise the spectra. As a clustering algorithm, hierarchical clustering analysis (HCA) with an appropriate distance function can be used in combination with other clustering algorithms, since there are studies that state the performance of HCA is better than k-means for clustering and segmenting the tissue regions based on histopathological evaluations [80]. In order to use the time efficiently and not to miss the parts of the scanned tissues during the experiments, an automated imaging procedure that starts with a low-resolution scan can be used. In this way, boundaries of tissue can be identified with an initial fast scan, and the information fed back for high-resolution scanning in the second or third run, depending on the reliability of the first result.

## Materials and methods

### Sample collection and preparation

Female BALB/c mice, six weeks of age, were sourced from the Animal Research Unit of Akdeniz University, Antalya, Turkey. A total of three subjects were involved in the study, in adherence to an experimental protocol approved by the Akdeniz University Faculty of Medicine’s Animal Care and Use Committee (Ethical Approval No. 2018.10.018). The mice were accommodated in a regulated environment with a light-dark cycle of 12 hours each and provided unrestricted access to food. To facilitate the formation of pregnancy groups, mature male mice of the same strain were utilized. The female mice were bred with these fertile males to instigate pregnancy, with the day of observed copulation plug serving as the first day of pregnancy. Placental samples were subsequently acquired on the 16*^th^* day of gestation. At the end of the study, the mice were humanely euthanized following ether anesthesia by means of cervical dislocation.

### Tissue processing and Hematoxylin and Eosin staining

The acquired placental tissues were immersed in 10% neutral buffered formalin for 24 hours to ensure thorough fixation. Upon completion of the fixation process, the tissues were thoroughly washed and dehydrated using a graded series of ethanol, subsequently cleared in xylene, and embedded in paraffin wax. Paraffin blocks were then prepared and sectioned at 5 *µm* thickness using a microtome. The obtained sections were adhered onto microscope slides and left to dry overnight at 37°C. These sections were then deparaffinized in xylene and rehydrated through descending concentrations of ethanol, prior to staining. Hematoxylin was first applied to stain the nuclear material, after which the sections were briefly washed and counterstained with Eosin, a cytoplasmic and extracellular matrix stain. Finally, after a series of dehydration steps in ascending concentrations of ethanol, the stained tissue sections were cleared in xylene and mounted with a resinous medium. Our H&E staining protocol provided clear and distinct images of cellular structures within the placental tissue, facilitating a thorough examination of the morphological changes that occurred at this developmental stage.

### Raman microspectroscopy and scanning

For the micro-Raman spectroscopy scans, a previously constructed lab-built Raman spectroscopy setup was employed [81]. A diode laser with 500 *mW* power and a wavelength of 785 *nm* was used. The laser beam was guided to the “Plan-Neofluar” 20x, 0.4 NA (Carl Zeiss Microscopy, LLC) objective lens using silver mirrors and a DMLP 805 dichroic mirror (Thorlabs). The beam width was expanded and collimated to fill the back aperture of the objective lens. The laser power at the back aperture of the objective was measured to be 57.65 *mW ±* 1.09 *mW*. The Raman scattered photons were collected using the same objective lens and directed into the same path but in the opposite direction of the laser input. These Raman scattered photons were then separated from the main line by a dichroic mirror. The separated Raman signal was filtered with Raman edge filters and coupled to a multimode fiber of NA 0.22 and connected to the QE Pro Raman spectrometer (Ocean Insight).

A motorized XY stage was utilized (Standa-8MTF-102LS05) and interconnected with MATLAB to scan the placental tissues. A common area for all samples was defined for the time efficiency of the scans. The samples were raster scanned in an area of 2.0 x 4.8 *mm* (101 x 241 pixels) with 20 *µm* steps and 300 *ms* integration time for each step. Thus, 24341 Raman spectra were collected from each mouse tissue. To denoise the spectra, a boxcar width of 2 pixels was used during the scans. Using these settings, placental tissues were scanned before and after deparaffinization to determine the effect of paraffin on Raman spectra and corresponding Raman maps. Raman spectra of an empty glass slide and a paraffin block were also collected to be used as background spectra and for contaminant identification, respectively, which are given in Fig S3 of the S1 File.

### Edge detection and preprocessing

The wavelength to Raman shift (*cm^−^*^1^) conversion was performed for the 785 *nm* laser wavelength. Since it is known that the region of the spectrum between 400 *cm^−^*^1^ and 665 *cm^−^*^1^ does not contain strong paraffin peaks, as validated both experimentally in Fig S3a of the S1 File and in the literature [40, 41], and the Raman intensity of the tissue was observed to be highest, this region was chosen for the edge detection procedure. These local minima were line-fitted and the fit was subtracted from the spectra as in the conventional linear baseline correction procedure. This generalized method was applied to both paraffinized and deparaffinized tissue spectra. In this way, the presence of paraffin did not affect the normalization of the spectra; hence, the edge detection performance for both the paraffinized and deparaffinized tissue spectra. After vector normalization was applied, the highest peak and expression of the tissue relative to the background was found to be at 510 *cm^−^*^1^. The tissue images at this wavenumber were used for edge detection of the tissue and masking of the non-tissue region. An example edge detection procedure for the tissue of mouse 1 is shown in Fig 5a. The user manually selects an area of background on the Raman map at 510 *cm^−^*^1^ as indicated by the yellow box in Fig 5a_1_. A threshold was set from the average value of the selected region and the tissue image was binarized once according to this threshold. All holes within the tissue region were filled using the built-in MATLAB function “imfill”. Then, the background mask was obtained by taking the inverse of the binary image. Finally, the average value of the masked region was used as a global threshold, which is less sensitive to user selection, and used to acquire the final binary image as depicted in Fig 5a_2_. The mask obtained from the inverse of the final binary image, shown in pink in Fig 5a_3_, was used to set the spectrum of the background pixels to Nan. Although 24341 different Raman spectra were collected from the placenta of each mouse, the number of spectra was reduced after edge detection. For the measurements before deparaffinization, the number of background eliminated tissue spectra were reduced to 16121, 14603, and 20280, whereas after deparaffinization, it was 15701, 13986, and 21460 for mice 1, 2, and 3, respectively.

**Fig 5.**
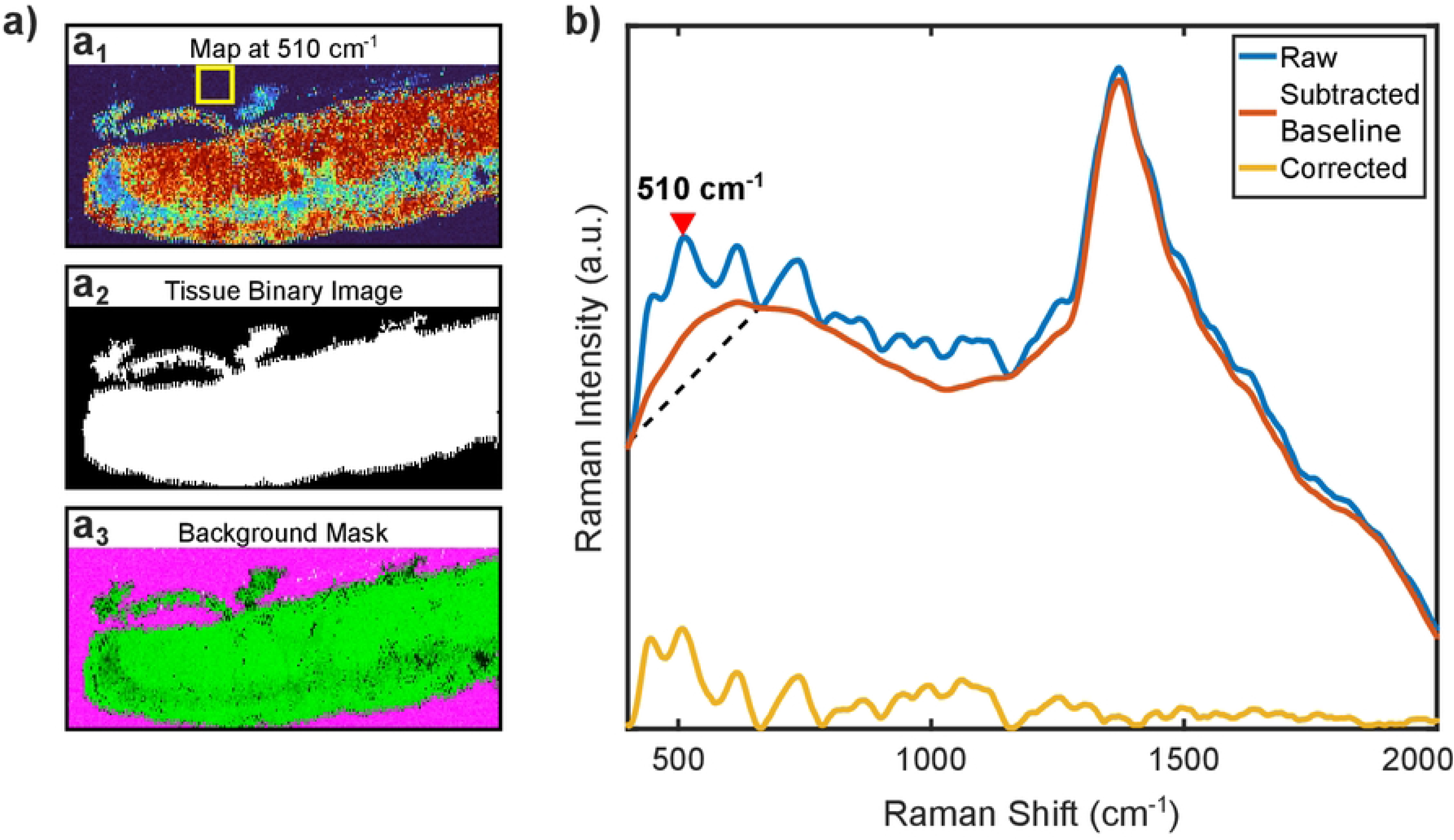
a) Edge detection flow. The yellow box in a_1_ represents the region selected by the user to determine a threshold for binarization. In a_2_, the final binarized image is displayed. Finally, in a_3_, the background mask and the tissue image are merged into a single image with pink and green colors, respectively. b) An example raw spectrum from a placenta tissue, subtracted baseline that includes modified polynomial fit with background subtraction of the glass substrate, and the corrected spectrum. The region between 400 *cm^−^*^1^ and 665 *cm^−^*^1^, and the peak at 510 *cm^−^*^1^ used for edge detection are also marked in the figure.

After edge detection, the spectra in the remaining pixels were cut between wavenumbers 400 *cm^−^*^1^ and 3000 *cm^−^*^1^ in the raw data. A modified polynomial baseline correction algorithm was then used for simultaneous baseline correction and glass background subtraction using a reference substrate spectrum [82, 83]. The glass spectrum was measured separately, given in Fig S3b of the S1 File, and used in the algorithm along with the fifth-order polynomial fitting. For deparaffinized tissues, the Raman spectra were cut between 400 *cm^−^*^1^ and 2000 *cm^−^*^1^, as this is the distinctive region for the biological samples [84]. Then, they were normalized using vector normalization. An example of a raw spectrum of placental tissue and the subtracted baseline that includes the glass spectrum as a background contaminant is shown in Fig 5b. The difference between the two gives the corrected spectrum. The figure also includes the region and Raman peak used for edge detection. However, when comparing the paraffinized and deparaffinized tissue spectra, the region between 2800 *cm^−^*^1^ and 3000 *cm^−^*^1^ was also added to the previous region and normalized together since there are two strong paraffin peaks here as seen in Fig 4b. The preprocessed spectral data and image data are uploaded to the open repository Zenodo [85].

### Machine learning and statistical analysis

As a data reduction step, PCA was applied to the pre-processed spectral data prior to classification. The acquired scores of each spectrum after PCA were sorted according to their TEV in decreasing order starting from the highest. Then, the TEVs were summed until a selected threshold of 95% was reached, and the scores to be used were determined as the number of summed TEVs. For each mouse, the number of scores used was reported in the supplementary table in Table S3.

The k-means clustering with a squared Euclidean distance algorithm was used to classify the spectral data. Selected scores of each mouse were fed separately into the k-means algorithm in MATLAB to label each spectrum. The number of clusters was chosen to be three, as a larger number of clusters did not result in further differentiation of meaningful zones or D and LZ. After k-means classification, the number of spectra corresponding to the number of pixels in the Raman clustering maps was found for each group as given in Table S3 of the S1 File.

As described in the text, group 3 was found to be due to the regions where the glass substrate was dominant; therefore, the statistical analyses focused only on the comparison between group 1 and group 2. For this purpose, automatically found peaks and their vicinity, limited to an interval of 4 *cm^−^*^1^, were examined for possible assignments given in the literature and selected accordingly [28]. The selected wavenumbers were marked as tentative assignments as shown in Fig 2b and Fig 2d. Their violin plots were then generated and plotted to compare the distribution of the data as shown in Fig 3. The violin plots were plotted using a modified function in MATLAB file exchange [86] at six different wavenumbers where the corresponding group colors are the same as in Fig 2. These six wavenumbers were chosen with respect to the diversity of macro assignments and the degree of visible differences in Raman intensities. For example, 1063 *cm^−^*^1^ was chosen for the lipid band, while 815 *cm^−^*^1^ was chosen for the proline and hydroxyproline bands. The median and mean values of the distributions are given within the violin plots for each group, and the vertical length of the blue boxes is called the interquartile range, which covers the 25*^th^* and 75*^th^* percentiles of the data within the groups. To quantitatively infer that these two groups differ from each other at the assigned wavenumbers within a certain significance, the built-in MATLAB function for the Wilcoxon rank sum test was applied to the tentatively assigned peaks. To control the family-wise error rate, the Bonferroni-Holm correction was applied to the p-values [87]. The multiple null hypothesis tests assuming that the two groups are equal were rejected at the 0.1% significance level since the values were calculated to be less than 0.001.

Before comparing the paraffin-embedded tissue images with the deparaffinized ones, an elimination was applied to the image data at the specified wavenumbers to remove cosmic rays and spectrometer overexposures. Therefore, a percentile elimination with a lower threshold of 0.1 % and an upper threshold of 99.9 % was applied to the data, i.e. the data outside the defined percentages were labeled as outliers. Since all tissue scans consisted of 24341 pixels, the elimination resulted in the conversion of 48 pixel values to Nan for each image. These pixels were replaced by the average spectrum of the remaining pixels prior to the spatial alignment of the images. A geometric transformation that aligns the paraffinized tissue image with the deparaffinized image at 510 *cm^−^*^1^ is found using the “imregtform” function in MATLAB. The first image was used as the reference image while the other one was used as the moving image. The resulting transformation was applied to all the paraffinized tissue images using the “imwarp” function. Non-overlapping pixels between the images that emerged after the transformation were cropped from all six images. All the obtained images were then compared to the deparaffinized tissue image at 510 *cm^−^*^1^ which was selected as the ground truth for the comparison. The process flow for the image alignment procedure is given for mouse 1 in Fig S5 of the S1 File. The images were then compared using the 2-D correlation coefficient function “corr2” in MATLAB. To avoid the Nan values assigned to the background regions in the edge detection step, the Raman images and their correlation coefficients were presented and calculated including the background data. However, the compared spectra were obtained only from the tissue region found in the edge detection step. After calculating the correlation coefficients between the images, a strength was assigned to these values. The correlation strength is defined in five steps from 0 to 1 with increments of 0.20, starting from very weak and ending at very strong, as given in the S1 File, Table S2.

### Histology and image analysis

High-resolution images of the stained sections were captured using a light microscope equipped with a digital camera (Zeiss, Oberkochen, Germany). These images facilitated a detailed examination of cellular and tissue-level changes within the placenta at E16. This enabled us to clearly identify and discriminate between the placental zones, each characterized by unique cellular arrangements and tissue structures. By enhancing the contrast between different cellular constituents and extracellular matrix, the H&E staining was particularly effective in demarcating the boundary lines of these zones. This detailed examination allowed us to generate precise topographical maps of the placenta and highlighted the inherent complexity and diversity within the placental tissue. Such accurate zone discrimination is vital to understand the intricate dynamics of placental development and function, and it provides a robust foundation for subsequent morphological and functional investigations.

### Visualization and programming

The visualizations were organized and created using Inkscape. Some of the illustrations in the graphical abstract were taken from SciDraw.io [88–90]. Others were either taken from free vector sites with a CC0 1.0 Universal (CC0 1.0) license or created in Inkscape.

MATLAB 2021b was used for all preprocessing and analysis. Algorithms and functions not found in MATLAB itself were obtained from the community in file exchange [86, 87] or written by the authors.

## Conclusions

In conclusion, three distinct zones of mature mouse placenta were biochemically and structurally distinguished using micro-Raman spectroscopy, which is a label-free technique. Clustering and spectral analysis of the collected data from three different mice revealed placental zones D, LZ, and JZ, with JZ being visually and biochemically distinct from the sum of the regions D and LZ. Tentative assignments and their relative distribution in the zones were shown to be consistent with H&E stained tissue images and the current literature. In addition, Raman imaging of paraffin-embedded placental tissues at specific wavenumbers was shown to be useful for easily accessing the morphology of the tissue without applying digital dewaxing algorithms, and the interfering effect of paraffin on tissue lipids demonstrated by image analysis.

## Supporting information

**S1 File. Supporting file that includes results of the other two mice placenta scans & information about the methods used in the analysis.**

